# The Role of Vision and Lateral Line Sensing for Schooling in Giant Danios (Devario Aequipinnatus)

**DOI:** 10.1101/2023.07.25.550510

**Authors:** Ben Tidswell, Annushka Veliko-Shapko, Eric Tytell

## Abstract

To protect themselves from predators, fishes often form schools with other fish. Previous work has identified abstract “rules” of schooling – attraction to neighbours that are far away, repulsion from neighbours that are too close, and alignment with neighbours at the correct distance – but we do not understand well how these rules emerge from the sensory physiology and behaviour of individual fish. In particular, fish use both vision and their lateral lines to sense other fish, but it is unclear how they integrate information from these sensory modalities to coordinate schooling behaviour. To address this question, we studied how the schooling of giant danios (Devario aequipinnatus) changes when they are either unable to see or unable to use their lateral lines. We found that giant danios were able to school normally without their lateral lines, but did not school in darkness. Surprisingly, giant danios in darkness had the same attraction and alignment properties as fish in light, potentially indicating that they do not feel as much risk in darkness. Overall, we suggest that differences among schooling species in sensory integration between vision and lateral line may depend on their natural predators and environment.

## Introduction

Schooling is a widely observed behaviour among many species of fish. Half of all fish species school at some point in their life, and a quarter school their entire lives (Shaw, 1978). Schooling allows fish to find food more easily and helps save energy while swimming (Ashraf et al., 2016; Reebs, 2000; Romenskyy et al., 2020), but its primary function is to help avoid predators (Herbert-Read et al., 2017; Ioannou et al., 2012; Jeschke & Tollrian, 2007; Parrish, 1989; Rieucau et al., 2016; Ward et al., 2011). By responding to the motion of their neighbours, fish in the school are able to gain information about the environment and pass that along to other members of the school – faster than they could if they swam on their own (Couzin, 2009; Gerlotto et al., 2006; Rieucau et al., 2016).

Early research on fish schooling used simple “rules of schooling” to model the choices that fish made that allowed the school to stay together (Aoki, 1982; Breder, 1954; Herbert-Read, 2016). In particular, three simple “rules” capture much of the behaviour of schools of fish and other animal aggregations: each animal is attracted to neighbours that are far away, repelled from neighbours that are too close, and tries to align with neighbours at the correct distance (Aoki, 1982; Herbert-Read, 2016). These early models assumed that fish had perfect knowledge of where the other fish in the school were, and they were able to pay attention to all the fish around them. But real fish must rely only on the information they are able to gather from their senses, which are limited to the few fish nearest to them (Jiang et al., 2017; Katz et al., 2011). Despite the importance and prevalence of schooling, it is still unclear how fish integrate information from their different senses to maintain a cohesive school. Therefore, the goal of the current study was to determine the roles of vision and the lateral line in schooling in giant danios (Devario aequipinnatus).

These two senses have long been thought to be the primary sensory mechanisms underlying fish schooling (Partridge & Pitcher, 1980). Originally, vision was posited as both necessary and sufficient for schooling (Breder, 1954). However, Partridge and Pitcher (1980) tested the roles of both the lateral line and vision in pollock and showed that they were able to school while blinded, which supports the idea that the lateral line is sufficient for schooling, at least in pollock. In contrast, similar research has found that many other fish species cannot school in darkness (Ginnaw et al., 2020; Kowalko et al., 2013; McKee et al., 2020; Patch et al., 2022; Ryer & Olla, 1998). This inability to school in darkness suggests that schooling with only the lateral line may not be a common ability among fish species, and that the degree to which fish rely on the lateral line or vision may vary greatly between species.

Moreover, the lateral line and visual systems work on different time scales and at different ranges, which means that together they provide more information than either one individually. The lateral line works over short ranges, but is processed rapidly by the nervous system (Bleckmann & Zelick, 2009), while the visual system has a longer range, but is processed more slowly (Pita et al., 2015). Of course, both of these systems can be compromised by environmental effect such has turbidity or turbulence, as well as injury or disease. Since these senses have their own limitations, and can be further limited by outside factors, the ability to rely on multiple senses would allow fish to maintain a cohesive school more reliably.

Previous work with giant danios has investigated how they school in flowing water and when one fish in a school has its lateral line ablated (Chicoli et al., 2014; Mekdara et al., 2018, 2019). Chicoli et al. (2014) found that giant danios in flowing water swam side by side more often than in still water. The school was also more polarized in flow, likely because the flow caused fish to orient upstream. Mekdara et al. (2018, 2019) also studied the role of the lateral line is schooling, but only ablating the lateral line of a single fish in order to see how its behaviour changed, instead of abating the lateral lines of all of the fish in the school. They found that a single fish could swim normally with a school immediately after its lateral line was ablated, but failed to school normally one to two weeks later even after its neuromasts had regenerated (Mekdara et al., 2018). In a follow up study they differentially ablated the anterior and posterior lateral lines, finding that without the anterior branch the fish could not match the velocity of nearby fish, without the posterior branch the giant danios had reduced tailbeat synchronization, and ablating any part of the lateral line caused fish to changed their formation relative to other fish (Mekdara et al., 2019).

The goal of this current study was to gain a better understanding of the role of vision and the lateral line in the schooling of giant danios when all fish in a school were affected by sensory limitations, examining how these senses contribute to this species’ ability to form a cohesive school and ultimately to avoid predators. We measured the three dimensional swimming kinematics of giant danios using high speed cameras. We limited vision by filming under infrared light with all visible lights turned off, and we limited lateral line sensing by ablating the sensory neuromasts with the aminoglycoside antibiotic gentamicin, which kills sensory hair cells in the neuromasts (Faucher et al., 2010; Mekdara et al., 2019). For each of these conditions we measured the nearest neighbour distance (NND), polarization, speed of the school, and turns made by the fish.

## Materials and methods

### Animals

For this set of experiments we used giant danios (Devario aequipinnatus) which were purchased through LiveAquaria (LiveAquaria, Rhinelander, WI) and from a PetSmart (PetSmart, Cambridge, MA). Before experiments, fish were kept in 40 litre aquarium tanks at 23°C and a conductivity of 400 μS, fed goldfish flakes daily (TetraFin, Blackburg, VA), and kept on a 12 hour night and day cycle. All experiments followed an approved Tufts University IACUC protocol (M2021-99 and M2018-103).

### Schooling Experimental Setup

For each of the schooling experiments, eight giant danios were randomly selected from the pool of forty available fish. Experiments were conducted in a Loligo Systems (Loligo Systems, Viborg, Denmark) flow tank, and fish were constrained to a filming area that was 60 cm long, 25 cm wide, and 25 cm tall and filled to a water depth of 20 cm.

On each day of filming, one of the three experimental sensory conditions was chosen: light, darkness, or lateral line ablation. For the experiments in darkness, lights were turned off in the room, light sources were covered, and filming was done using 850 nm IR lights (CMVision, Houston, TX). The spectral sensitivity for vision in near infrared in giant danios is not known, but their visual sensitivity is at between 50 and 150 times lower at 750 nm than at 500 nm (van Roessel, 1997), suggesting that they have even lower sensitivity at 850 nm. Similarly, closely related zebrafish (Brachydanio rerio) have a substantial drop in optomotor responsiveness at 840 nm (Matsuo et al., 2021), suggesting that zebrafish, and by extension giant danios, have much reduced vision at wavelengths greater than 840 nm. For the experiments with lateral line ablation, the giant danios were placed into gentamicin treated water (0.001% for 24 h, 5 litres, Sigma-Aldrich, St Louis, MO) following previously established protocols (Mekdara et al., 2018). Half of the recorded trials were in flow (at a speed of two body lengths, BL, per second; 2 BL/s) and half were in still water, with these assigned in a random order in blocks of three trials. The flow trials allowed us to see if the patterns in fish behaviour were consistent when the fish were oriented towards flow, which is particularly relevant for giant danios since they natively live in flowing water (Dey et al., 2014)

During each of these experiments, fish were filmed for thirty five second at 60 frames per second (fps) using two high speed cameras (Phantom M320, Vision Research, Wayne, NJ). We used two cameras in order to allow for the 3D reconstruction of the position of the fish as our setup was deep enough that the height of the school could have cause important projection errors (Supplemental Materials). These two cameras were synchronized, which allowed us to reconstruct 3D trajectories of all the fish in the school. The cameras were calibrated using open source software DLTdv8 and the easyWand program (Hedrick, 2008; Theriault et al., 2014). We used the open source software Deeplabcut (Lauer et al., 2021; Mathis et al., 2018; Nath et al., 2019) to train neural networks to track fish in each individual video, and were then triangulated with camera calibrations using DLT code written in Python to compute 3D positions (Python Software Foundation, 3.9.12).

### Analysis of Schooling Behaviour

With the 3D coordinates of all of our fish, we calculated the kinematics of the fish school. To follow the standards set by other papers (Couzin et al., 2002; McKee et al., 2020; Mekdara et al., 2018; Partridge & Pitcher, 1980), we used the 3D coordinates of fish bodies in order to calculate the kinematics of the fish school as defined by the mean nearest neighbour distance across all fish in the school 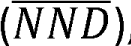, the overall speed of the school, and the mean polarization of the school. These values were calculated as follows. Nearest neighbour distance in 3D is

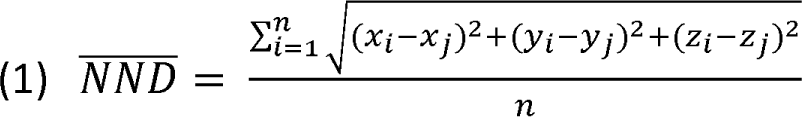

where (x_i_,y_i_,,_i_) represents the snout position of a particular fish, ; is an index for a particular fish, 1 is the index for the closest fish to fish ;, and n is the total number of fish. Polarization is

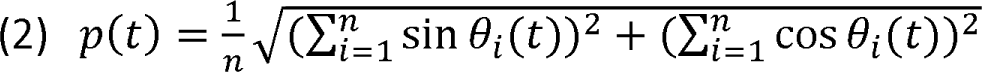

where θ_i_ is the heading angle of fish ; (Couzin et al., 2002; Jolles et al., 2017). The overall speed of the school is

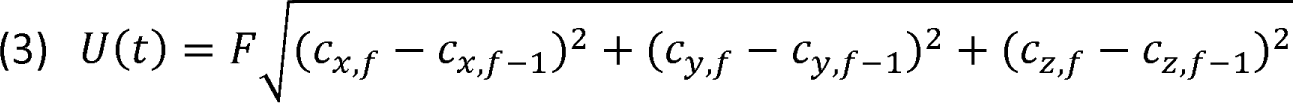

where *c_j_* = (*c_x,f_, c_y,f_ c_z,f_*) is the point representing the center of the school in frame *f*, as calculated by taking the average X, Y, and Z coordinates of all the fish in the school, and F is the frame rate (60 FPS). Note that all of these metrics depend on accurately calibrated 3D positions. Two-dimensional data from a single camera can introduce substantial error due to parallax (Fig. S1).

To calculate the number of subunits in the school, we first created a distance matrix between all the fish in the school. We then converted that into a matrix of zeroes and ones, where an element *G_ij_* was a one if fish ; was within two body lengths of fish 1, and a zero if they were not. This matrix was used to create a node and edge graph connecting the fish that were within two body lengths of each other. We then calculated the number of completely separate subnetworks made by these connections using the python package NetworkX (Hagberg et al., 2008). We used the number of subnetworks as the number of different subunits that the school had formed. Calculating the number of subunits in the school allowed us to get another sense of the way fish were schooling, as NND can obscure the formation of multiple subunits if the fish stay close enough to the other fish they have grouped up with (Figure 1).

**Figure 1.**
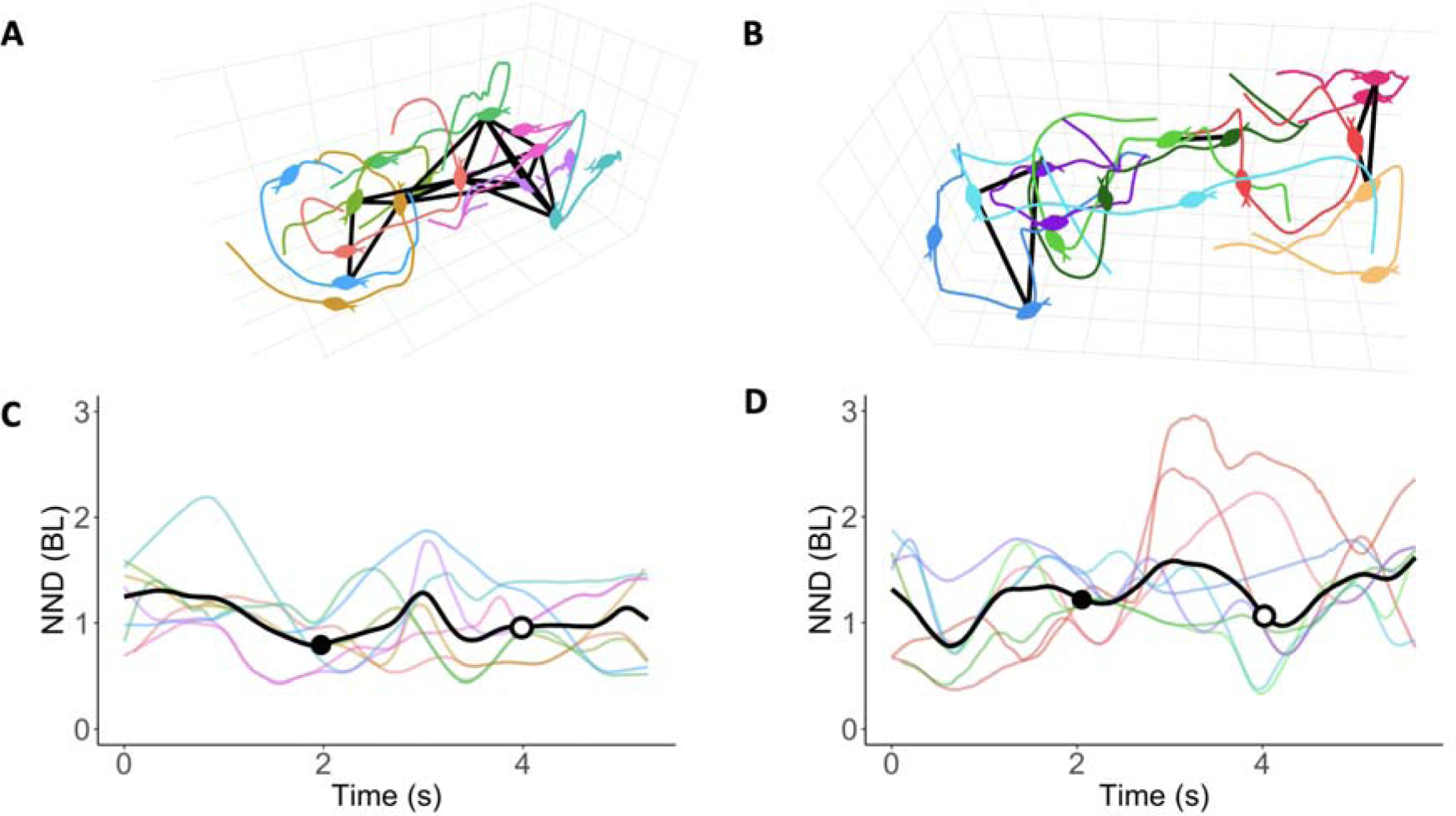
Kinematics measurements of nearest-neighbour distance and subunits in schooling giant danios. The path of each fish in 3D space is shown here for a specific trial in light (A) and darkness (B). Individuals who are within 2 body lengths of each other at the 2 second time point are connected by black lines, and considered part of the same schooling subunit. The location of each fish at the 4 second timepoint are also included, but their connections are not shown. The nearest-neighbour distance for each fish in light (C) and darkness (D) are shown, along with the mean among all members (black curve). The NND at 2 and 4 seconds are specifically highlighted with a closed and open dot, respectively.

### Turning behaviour

Turns were identified by taking the dot product between the heading unit vector of each fish, and the heading vector twenty frames afterwards. These dot products were averaged over every ten frames. We identified any peaks that had a change of heading of greater than 45°, which were then considered the point in time where the turn occurred. We counted the number of fish to the left and right of the turning fish before the turn, based on their location in the flat 2D plane.

### Null model for turning

To verify that any effects seen on turning based on sensory conditions were based on changes to fish’s schooling behaviour, not because of the physical constraints of the tank, we constructed a null model for fish turning behaviour in our tank. To that end we collected video of trials where we only placed one fish in the tank. This was done under all sensory conditions, but only in still water. We then used Deeplabcut to track these individual fish, and then combined eight of these tracks to create an artificial “school”. Because we created a “school” of eight fish by combining the tracked points of fish that were filmed at different times, there would be no way for them to turn based on the movements of the other fish in their “school”. The probability of making a right turn based on the number of fish on their side was compared in each of the different sensory conditions in order to validate that the real schools behaved differently than the artificial ones.

### Statistical Analysis

For all of the statistical analysis preformed, all measurements of the schooling kinematics were first averaged over individual tailbeats, because fish change their behaviour at the time scale of a tailbeat. To this end we found the average tailbeat duration within all the trials, and then grouped the measurements of fish schooling over windows of that duration and averaged each measurement within that window. Thus, each datapoint is represented as the average behaviour of fish during one tailbeat, which acts as a biologically relevant unit of time for these giant danios. However, because these points follow one another in time, they are autocorrelated. To reduce the effects of autocorrelation we analysed every third tailbeat in our data analysis, which we found to have the largest reduction in autocorrelation while still preserving the most data (Fig. S1).

All of our statistical analysis was done in Python (Python Software Foundation, 3.7.12) or R (Version 4.2.2) We made use of the statistical packages lme4 for running generalized linear mixed effects models (GLMMs) (Bates et al., 2015) and car for analysing these models (Weisberg S, 2019). The NND, polarization, school speed, and probability of turns were all tested using GLMMs in order to determine if there were significant differences among the different treatment groups, and to account for variation caused by using the same school of fish for multiple trials on the same day. The darkness and ablation conditions were each compared to the light and intact lateral line condition to determine if lacking access to a sense caused a significant change in aspects of schooling behaviour. All statistical tests were considered significant at P < 0.05.

## Results

Figure 2 shows the changes in the school in the dark and when all of the fish had their lateral lines ablated. In still water, fish in darkness swam significantly further from their neighbours (Fig. 2A and Table 1; F(1,81) = 6.875, p < 0.001), but with the same polarization (Fig. 2B and Table 1; p = 0.89), and average speed of the school (Fig. 2C and Table 1; p = 0.98). Importantly, they were split up into significantly more schooling subunits (Fig. 2D and Table 1; p = 0.003) compared to fish in light. While polarity and speed were the same for fish in darkness, the fact that they broke up into many more subunits and had a higher average NND shows that they were not one cohesive school. In flowing water, the fish in darkness behaved statistically similarly to fish in light: there were no significant differences in any of our metrics in flowing water between the control schools and schools in the dark (Fig. 2, right column and Table 1; p > 0.3 in all cases).

**Figure 2:**
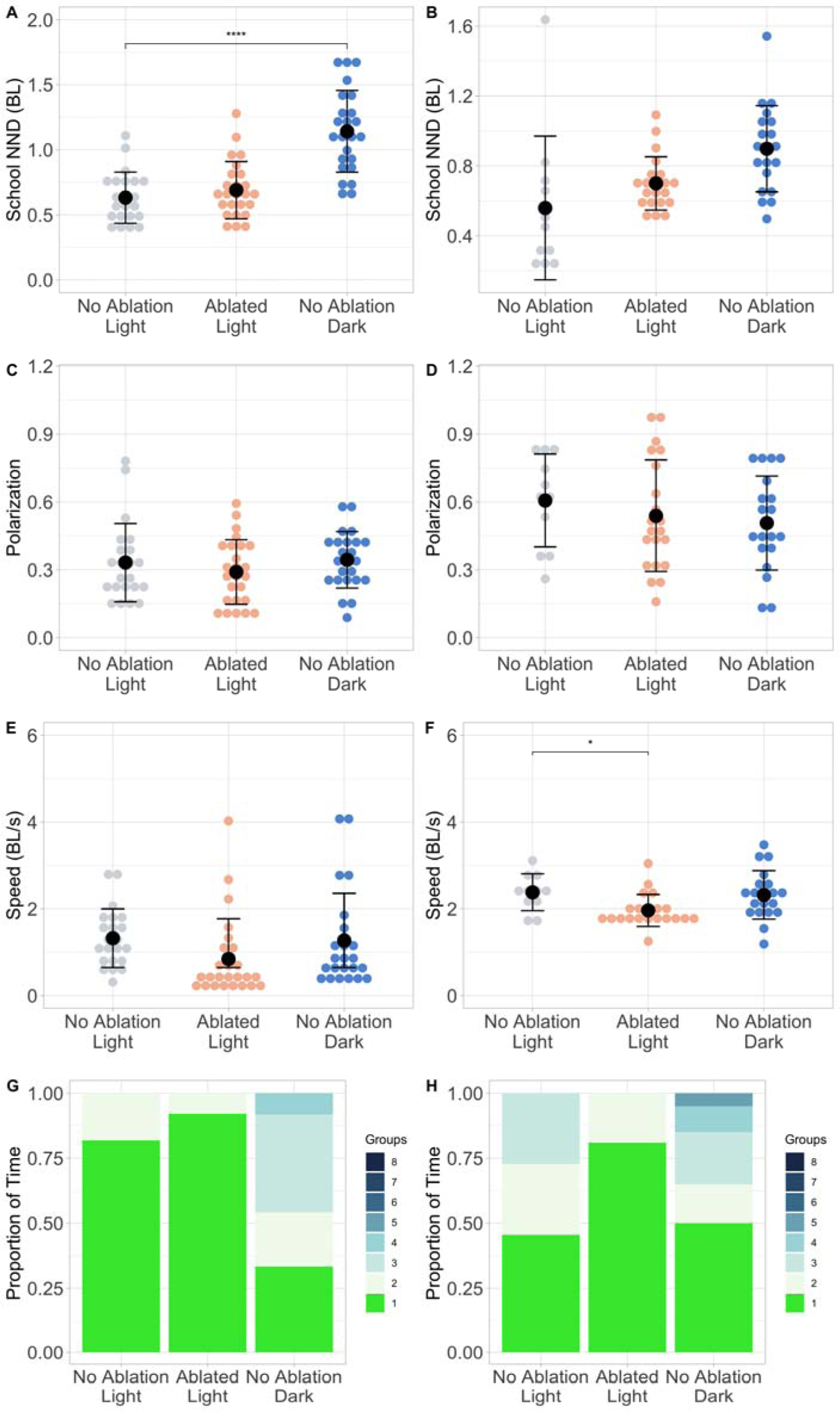
Giant danios form cohesive schools without their lateral lines, but school further from one another and form more schooling subunits in the dark. The NND (A, B), polarization (C, D), speed of the school (E, F), and number of subunits in the school (G, H) was measured in light, with lateral lines ablated, and in darkness. All plots on the left side (A, C, E, G) are in still water, while the plots on the right side (B, D, F, H) are in water flowing at 2 BL/s. Significant differences between sensory conditions are indicated by the asterisks, and were found by estimated marginal means with post hoc tests (* P<0.05, ** P<0.01, *** P<0.001, **** P<0.0001).

**Table 1:** Results of the GLMMs testing for the effect of darkness or ablation on the schooling behaviour of giant danios. P values show differences relative to the control condition. Significant effects are shown in bold.

| Measurement | Still or Flowing Water | Sensory Condition | DF | X<sup>2</sup> | P-Value |
| --- | --- | --- | --- | --- | --- |
| NND | Still | Ablation | 68 | 0.19 | 0.66 |
| **NND** | **Still** | **Darkness** | **68** | **19.29** | **< 0.001** |
| NND | Flowing | Ablation | 49 | 0.00 | 0.99 |
| NND | Flowing | Darkness | 49 | 0.86 | 0.35 |
| Polarization | Still | Ablation | 68 | 0.52 | 0.47 |
| Polarization | Still | Darkness | 68 | 0.02 | 0.89 |
| Polarization | Flowing | Ablation | 49 | 0.02 | 0.89 |
| Polarization | Flowing | Darkness | 49 | 0.24 | 0.63 |
| Speed | Still | Ablation | 68 | 0.82 | 0.36 |
| Speed | Still | Darkness | 68 | 0.00 | 0.98 |
| **Speed** | **Flowing** | **Ablation** | **49** | **5.87** | **0.02** |
| Speed | Flowing | Darkness | 49 | 0.12 | 0.73 |
| Number of Subunits | Still | Ablation | 68 | 0.07 | 0.79 |
| **Number of Subunits** | **Still** | **Darkness** | **68** | **8.94** | **0.003** |
| Number of Subunits | Flowing | Ablation | 49 | 0.81 | 0.37 |
| Number of Subunits | Flowing | Darkness | 49 | 0.82 | 0.36 |

Ablating the lateral line of the fish with gentamicin produced relatively few statistical changes in the schooling kinematics of our fish. In still water, schools of treated fish showed no significant differences with control fish (Table 1; p > 0.3 in all cases) for nearest neighbour distance (Fig. 2A), polarization (Fig. 2B), speed of the school (Fig. 2C), and number of schooling subunits (Fig. 2D). In flowing water, the only significant difference was for the average speed of the school, which was significantly lower in treated fish than in control (Fig. 2C and Table 1; p = 0.02). While the speed of the treated school was different in flowing water, ablated fish performed almost exactly the same as fish with their lateral lines intact, indicating that treated fish were still able to maintain a cohesive school.

### Turning Behaviour

Turning behaviour in giant danios does not differ significantly across sensory conditions. Across all conditions, we found that fish were significantly more likely to turn towards where there were more fish, and that the likelihood of turning towards the majority of fish was similar even when fish had an ablated lateral line or were swimming in darkness. Overall, we found that the turns fish made in each experimental condition were influenced by the number of fish on either side, in both still (Fig. 3A and Table 2; p < 0.001) and flowing water (Fig. 3B and Table 2; p < 0.001). In still water, we found that there was not a significant change in the turning behaviour of fish in darkness (Table 2; p = 0.17) or with an ablated lateral line (Table 2; p = 0.92), compared to their behaviour in the light. In flowing water, there was not a significant change from control to swimming in darkness (Table 2; p = 0.23), but fish with ablated lateral lines were less influenced by the number of fish on either side of them (Table 2; p = 0.01).

**Figure 3:**
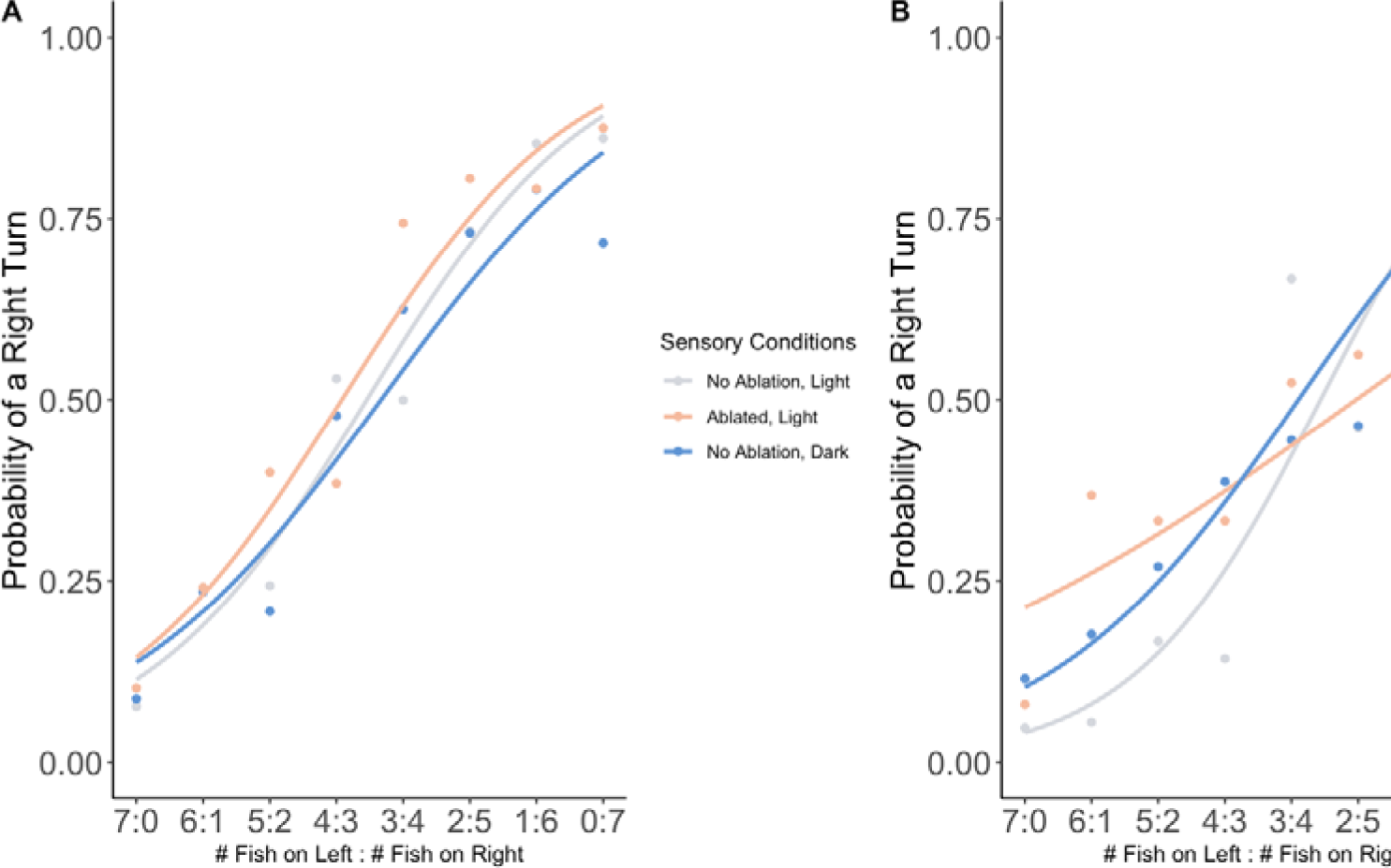
In all sensory conditions, fish are more likely to turn toward larger congregations of fish. The probability of turning towards the right, as a function of the number of fish to on the left and right (left:right) of the focal fish. (A) The probability of making a right turn in still water. (B) The probability of making a right turn in water flowing at 2 BL/s

**Table 2:** Results of the GLMMs testing for the effect of darkness or ablation on the turning behaviour of Giant Danios, with respect to the number of fish on either side of them (FNR)

| Still or Flowing Water | Factor | DF | $\chi^2$ | P-Value |
| --- | --- | --- | --- | --- |
| **Still** | **FNR** | **1043** | **398.05** | **< 0.001** |
| Still | Ablation | 1043 | 1.06 | 0.30 |
| Still | Darkness | 1043 | 0.79 | 0.37 |
| Still | FNR:Ablation | 1043 | 0.01 | 0.92 |
| Still | FNR:Darkness | 1043 | 1.91 | 0.17 |
| **Flowing** | **FNR** | **583** | **170.51** | **< 0.001** |
| Flowing | Ablation | 583 | 0.96 | 0.33 |
| Flowing | Darkness | 583 | 2.56 | 0.11 |
| **Flowing** | **FNR:Ablation** | **583** | **6.42** | **0.01** |
| Flowing | FNR:Darkness | 583 | 1.43 | 0.23 |

The similarity across sensory conditions is not an artifact of the size or shape of our filming arena. We compared the turning behaviour of a school of fish to an artificial “school” (the null model constructed by overlaying eight tracks of a single fish swimming in our tank). In each case, the real school behaved differently than the artificial one. Specifically, the probability of making a right turn, based on the ratio of fish on the left and right of the focal fish (Fish Number Ratio or FNR in statistical tests), was different when they were schooling in light with their lateral line intact (Fig. S3 and Table S1; p = 0.045), when their lateral line was ablated (Fig. S3B and Table S1; p = > 0.001), and when the fish were in darkness (Fig. S3C and Table S1; p = 0.024). Therefore, the effects we saw were not because of the constraints of our tank, but because of actual changes in fish schooling behaviour.

## Discussion

Our experiment addressed the role of the lateral line and vision in schooling behaviour. We found that giant danios do not form a cohesive school in darkness, as they had a much higher nearest neighbour distance and formed more schooling subunits than fish in light, indicating that the school was much more spread out and broken apart. In contrast, fish that had their lateral lines ablated were still able to school, maintaining cohesive groupings, though they were a bit more spread out than fish with their lateral lines intact. While these results were not the same when the fish were in flowing water, this may have been due to the influence of the flow as an outside orienting force. The ability of giant danios to school without their lateral lines but not in darkness suggests that, for this species, vision is necessary and sufficient for schooling.

These results provide us with more information about the sensory systems that fish rely on for schooling. Our finding that fish can school without their lateral lines comes into a divided literature on schooling and the lateral line. Previous work by Partridge and Pitcher (1980) and Mekdara (2018) found that saithe and giant danios could school without their lateral lines, but Faucher (2010) and McKee (2020) both found that rummy nosed tetras were unable to school properly without their lateral lines, and Middlemiss (2017) found similar results for yellow-eyed mullet. This indicates that while the lateral line may be useful for schooling, it may not be required for all fishes.

Our work adds more confirmation to the literature that most fish species cannot school in complete darkness. While Partridge and Pitcher (1980) found that an individual saithe could be part of a larger school without vision, but Kowalko (2013), Ginnaw (2020), and McKee (2020) respectively found that entire schools of Mexican tetras, three spined sticklebacks, and rummy nosed tetras were unable to school in darkness. Older work by Hunter (1985), Higgs and Fuiman (1996), and Miyazaki (2000) also all found that there was a level of darkness light beyond which fish would no longer form a school. Our study also found that fish could not school in darkness.

Even though giant danios do not form a cohesive schooling group in darkness, they still clearly sense and respond to one another. In darkness, they turn towards their neighbours just as often as they do in the light (Fig. 3), a form of the attraction “rule”. They also stay well-aligned with their neighbours (Fig. 4) and the school stays polarized (Fig. 2C,D). These results suggest that giant danios may be capable of schooling in darkness, using only their lateral lines to sense one another; instead, they may be choosing not to school. We hypothesize that they feel less risk in the darkness, since they are less exposed to visual predators, and therefore, they do not feel the pressure to form a cohesive school.

Other fish species also change behaviour based on their perception of risk (Rodriguez-Pinto et al., 2020; Ryer & Olla, 1998). For example, herring school more tightly when over a net that contrasted the colour of their bodies compared to a looser school over a net that was closer to their body colour (Rieucau et al., 2016). Golden shiners also school more closely when exposed to an alarm substance called Schreckstoff, released when neighbours are injured. Schreckstoff is thought to increase perceived risk, and leads to a tighter school (Sosna et al., 2019). In contrast, guppies Poecilia reticulata form shoals when threatened by a predator in clear water, but do not cohere well in turbid water, even with the threat of a predator (Kimbell & Morrell, 2015b). Kimbell and Morrell (2015b) suggest that the lack of cohesive schools in turbid water is not because guppies do not perceive the risk, but rather that they are not capable of shoaling cohesively without vision.

More broadly, differences in school behaviour among species may be explained by ecological or environmental differences in the habitats the species occupy. Turbid water can change the escape behaviours of schooling species, which adjust their strategies to account for a lack of vision (Kimbell & Morrell, 2015a). Previous studies have also show that environmental factors can change the number of neuromasts for subpopulations within a species (Fischer et al., 2013; Kelley et al., 2017; Spiller et al., 2017). It is therefore not unreasonable to consider that fish living in clearer water may depend more on vision than the lateral line, and those that live in turbid water or that school at night may adapt to depend more on the lateral line. However, this is an area that needs further study, and is complicated by the fact that many schooling fish are not caught naturally, but are purchased from breeders. Perception of risk may also depend on habitat and ecological interactions: fish that are preyed on more by visual predators like birds may perceive less risk in the dark than those that are preyed on by other fishes (Rieucau et al., 2016). Thus, differences between the environments where fish species evolved to live may also make a large difference in how much they rely on different sensory systems. While extreme environmental differences are known to have caused extreme sensory changes to blind cavefish (Kowalko et al., 2013; Kulpa et al., 2015; Patch et al., 2022), more subtle changes to the environment may still have profound impacts.

For example, McKee et al. (2020) found that rummy-nosed tetras Hemigrammus rhodostomus could not school in the dark, but also schooled differently without their lateral lines. In contrast, in our study, giant danios showed no significant changes in schooling behaviour without the lateral line. Rummy-nosed tetras tend to live in murky and slow moving water where visibility is low (Lewis et al., 2000). Meanwhile, giant danios generally are found in clear streams where there may be more visibility, but also more interference when trying to sense other fish due to the natural flow and turbulence of rivers (Dey et al., 2014). Tetras may need their lateral lines more since they often cannot see their neighbours as well, while Giant Danios rely more on vision because they are usually able to see the other fish in the school around them. It seems likely that the different life histories of these fish alter their reliance on each of their senses, thus changing how the school responds when those senses are removed.

From the work we have done here, it is clear that further work should be done to systematically compare the schooling behaviour of many different fish species from a variety of environments and with a variety of life histories to understand what environmental pressures may change fishes’ reliance on vision or the lateral line. It is also important to be mindful of how stress and the laboratory or testing environment can affect fish schooling and how this interacts with the way fish use their senses to school.

## Conclusion

In this study, we examined how vision and flow sensing impact the schooling behaviour of giant danios. Our results showed that vision is crucial for maintaining a cohesive school, while the lateral line system is not. Interestingly, we also observed that fish that were unable to see were still able to turn towards the majority of the school, suggesting that the lateral line sense on its own may provide relevant information about the location of other fish In darkness. However, our study still clearly shows that vision is necessary and sufficient for giant danios to school.

## Supplemental Material

**Supplemental Figure 1:**
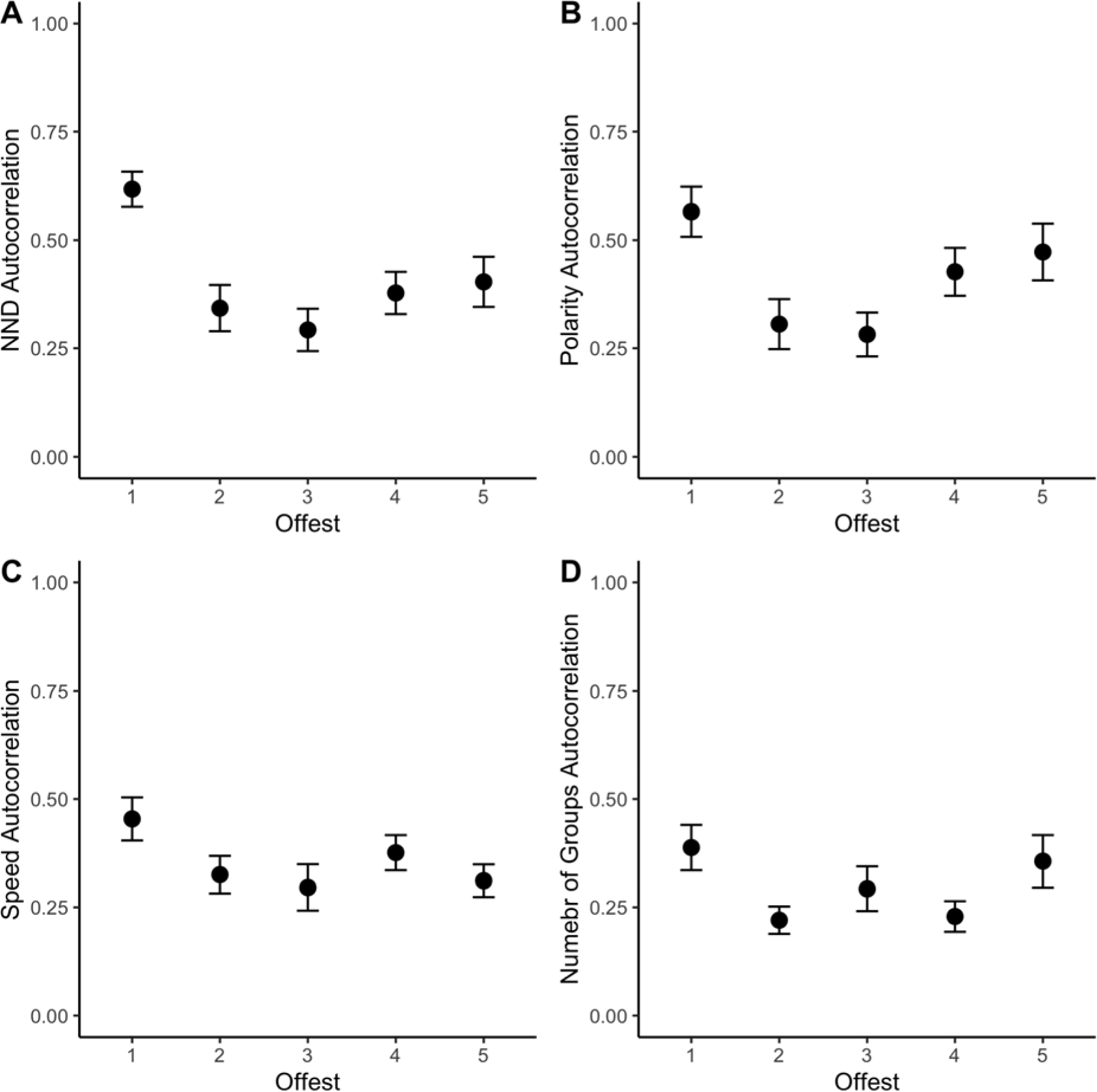
A lag of three provides the overall largest decrease in autocorrelation while still retaining the most data. The autocorrelation of (A) NND, (B) polarity, (C) speed, and (D) the number of schooling subunits is shown here for a given set of offset values of the data. We sound that there was the overall the largest decrease in autocorrelation between datapoints from each tailbeat with an offset of three. While greater offsets had the same or slightly decreased autocorrelation, an offset of three also preserves more of the data.

**Supplemental Figure 2:**
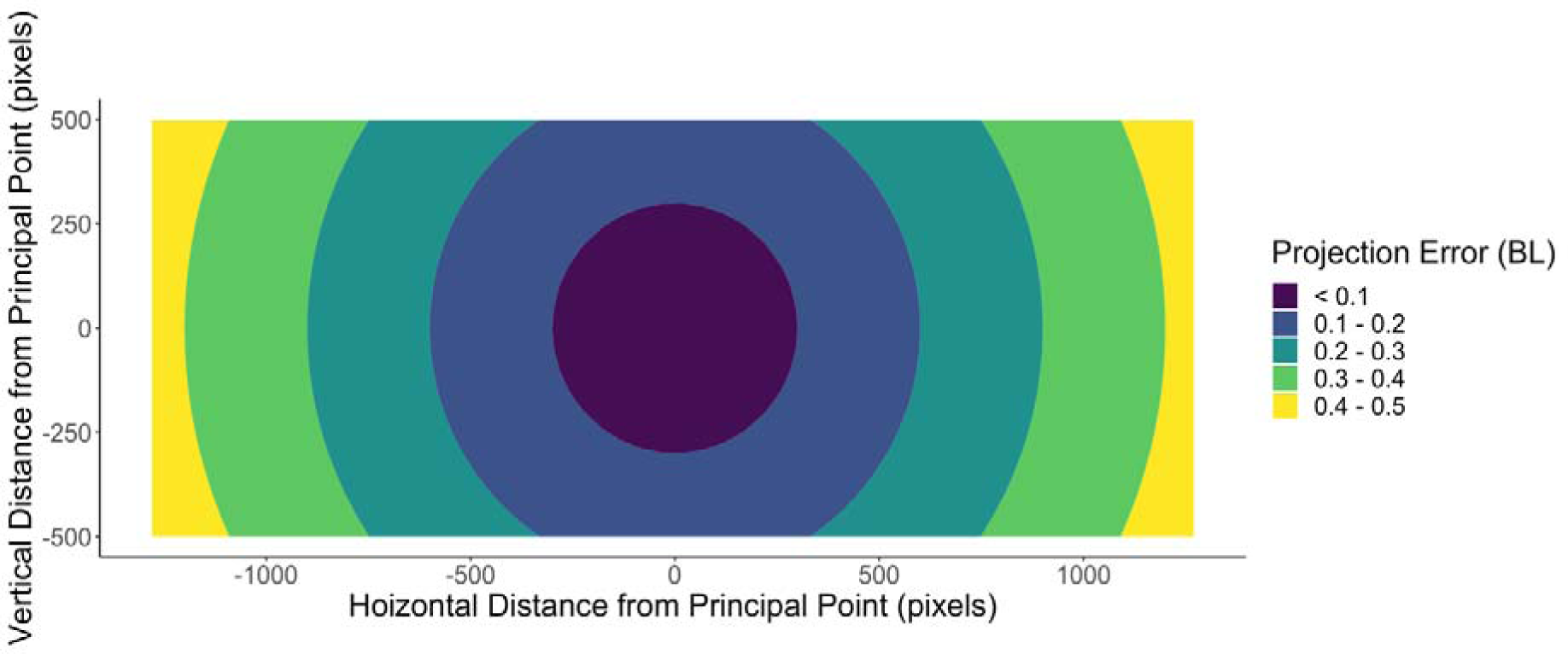
Using 3D data prevents up to half a bodylength of projection error in our setup The estimated projection error based on our camera size, distance from the school, and average height of the school (1 BL).

**Supplemental Figure 3.**
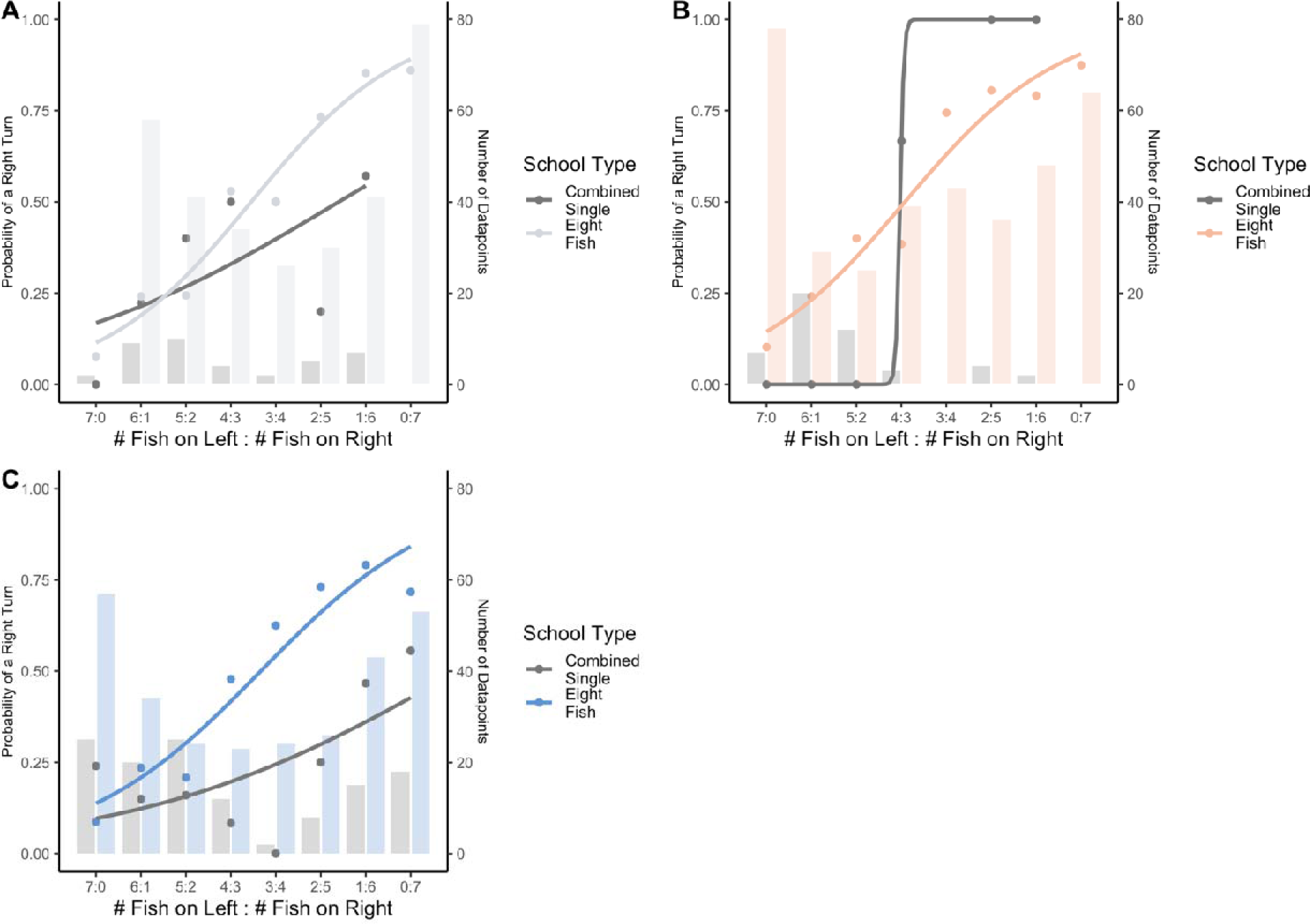
Turning behaviour depends on the numbers of neighbouring fish, not the constraints of the tank. The probability of turning towards the right, as a function of the number of fish to on the left and right (left:right) of the focal fish, comparing real schools to the artificial school (the null model). Each panel shows a different sensory condition: (A) when fish had their lateral line and were in the light, (B) when fish had their lateral line ablated, and (C) when fish were in darkness. The bar graphs show the number of turns that correspond to each data point. Note that the single fish “schools” showed far fewer turns overall than the schools of eight fish for the same number of trials.

**Supplemental Table 1:** Results of the GLMMs testing for the differences in the turning behaviour of artificial “schools” and real schools of eight fish.

| Condition | Factors | DF | X<sup>2</sup> | P-Value |
| --- | --- | --- | --- | --- |
| Light | SchoolType | 506 | 1.07 | 0.301 |
| **Light** | **FNR:SchoolType** | **506** | **4.02** | **0.045** |
| **Ablated** | **SchoolType** | **500** | **5.43** | **0.02** |
| **Ablated** | **FNR:SchoolType** | **500** | **17.46** | **< 0.001** |
| **Darkness** | **SchoolType** | **638** | **11.78** | **0.001** |
| **Darkness** | **FNR:SchoolType** | **638** | **6.07** | **0.024** |

